# Optimized phylogenetic clustering of HIV-1 sequence data for public health applications

**DOI:** 10.1101/2022.01.14.476062

**Authors:** Connor Chato, Yi Feng, Yuhua Ruan, Hui Xing, Joshua Herbeck, Marcia Kalish, Art F. Y. Poon

## Abstract

Clusters of genetically similar infections suggest rapid transmission and may indicate priorities for public health action or reveal underlying epidemiological processes. However, clusters often require user-defined thresholds and are sensitive to non-epidemiological factors, such as non-random sampling. Consequently the ideal threshold for public health applications varies substantially across settings. Here, we show a method which selects optimal thresholds for phylogenetic (subset tree) clustering based on population. We evaluated this method on HIV-1 *pol* datasets (*n* = 14,221 sequences) from four sites in USA (Tennessee, Seattle), Canada (Northern Alberta) and China (Beijing). Clusters were defined by tips descending from an ancestral node (with a minimum bootstrap support of 95%) through a series of branches, each with a length below a given threshold. Next, we used *pplacer* to graft new cases to the fixed tree by maximum likelihood. We evaluated the effect of varying branch-length thresholds on cluster growth as a count outcome by fitting two Poisson regression models: a null model that predicts growth from cluster size, and an alternative model that includes mean collection date as an additional covariate. The alternative model was favoured by AIC across most thresholds, with optimal (greatest difference in AIC) thresholds ranging 0.007–0.013 across sites. The range of optimal thresholds was more variable when re-sampling 80% of the data by location (IQR 0.008 – 0.016, *n* = 100 replicates). Our results use prospective phylogenetic cluster growth and suggest that there is more variation in effective thresholds for public health than those typically used in clustering studies.

## INTRODUCTION

Identifying clusters of infections with shared characteristics that imply a common origin is a fun-damental goal for epidemiological research and surveillance. For example, the ongoing global SARS-CoV-2 pandemic was first detected as a cluster of cases of unexplained viral pneumonia [1]. Clusters continue to identify outbreaks of SARS-CoV-2 transmission, informing lockdown protocols and identifying subpopulations for prioritized allocation of public health resources [2, 3]. Conventionally, clusters are identified by infections that are sampled within a relatively short time frame (temporal clustering), in association with a defined space (spatial clustering), or both. However, molecular sequence data have been used increasingly to cluster infections by their genetic similarity, which can provide a complementary or surrogate measure of their spatial or temporal proximity. There are now many examples of genetic clustering applied to RNA viruses, including Ebola virus [4, 5], human immunodeficiency virus type 1 (HIV-1) [6–8], hepatitis C virus [9], and coronaviruses associated with Middle Eastern respiratory syndrome (MERS-CoV) [10] and the 2003 outbreak of severe acute respiratory syndrome (SARS-CoV) [11]. The rapid evolution of many RNA viruses favours the use of genetic clustering because mutational differences can accumulate between infections in a matter of weeks or months [12, 13]. When transmission occurs on a similar time scale, a molecular phylogeny reconstructed from infections of an RNA virus will be shaped in part by its transmission history [14]. In this context, a phylogeny is a tree-based model of how infections are descended from their common ancestors.

HIV-1 has been particularly targeted for applications of genetic clustering. This is driven not only by HIV-1’s rapid evolution and global impact, but also by the availability of large sequence databases in many clinical settings from routine screening for drug resistance mutations. For instance, continual updates to sequence databases make it feasible to monitor the emergence and growth of genetic clusters over time [6, 15, 16]. Clustering methods have been used retrospectively to characterize populations associated with elevated rates of HIV-1 infection [8, 17–20], to identify potentially linked co-infections of HIV and hepatitis C virus [9, 21], superinfection by multiple HIV-1 subtypes [22] and transmitted drug resistance [23]. Clustering has also been proposed to support HIV-specific prevention methods such as pre-exposure prophylaxis (PrEP), which require a precise understanding of high-risk populations to optimize the distribution of public health resources [24, 25]. Finally, the lack of an effective vaccine demands a continuous assessment of priority populations for testing and antiretroviral treatment [26].

Over the last decade, there has been a growing diversity of genetic clustering methods, many of which were specifically designed and validated on HIV-1 sequence data [7, 27–32]. These methods can be broadly categorized by whether or not they require the reconstruction of a molecular phylogeny [33]. For example, Cluster Picker [27] is currently among the most widely-used genetic clustering methods for HIV-1 based on numbers of citations in the literature. Cluster Picker defines each cluster as a subtree in the phylogeny. A subtree is a portion of the phylogenetic tree that consists of an ancestral node and all of its descendants; in evolutionary terminology, a subtree is a monophyletic group. Cluster Picker searches for subtrees where (1) the total length of branches between any pair of tips in the subtree (*d*) is always below a threshold *d*_max_, and (2) the bootstrap support associated with the ancestral node exceeds a threshold *b*_min_. A bootstrap support value (b) is a measure of reproducibility — how often we expect to reconstruct a node ancestral to exactly the same set of tip labels from a hypothetical new data set of equivalent dimensions to the original data [34]. In practice, *d*_max_ and *b*_min_ are typically set to values in the range of 0.01 to 0.05 expected nucleotide substitutions per site and 90% to 95%, respectively [20, 27, 35, 36].

Clustering methods that do not reconstruct a phylogeny are also in widespread use for HIV-1 [16, 37–39]. For example, HIV-TRACE [7] employs a genetic distance [40] that reduces two sequences to a number quantifying the extent of their evolutionary divergence. Clusters are assembled from pairs of sequences whose distances fall below some predefined threshold. The advantages of distance-based clustering is that pairwise distances are rapid to compute and yield immutable quantities; these distances do not change with the addition of sequences to the database. In contrast, reconstructing a phylogeny with additional sequence data can change the branch lengths and bootstrap support values associated with previously-defined clusters [36]. Consequently, the use of clustering for continuous monitoring of an HIV-1 sequence database (*i.e*., to track the growth of clusters) has tended to focus on distance clustering methods [16, 39]. Tracking cluster growth can provide more informative indicators for public health decisions. For instance, large clusters tend to emphasize historical outbreaks that are no longer active [39, 41].

Nevertheless, phylogenetic clustering remains more prevalent in the infectious disease literature [42]. Clusters generated from pairwise distances tend to have a high density of connections (edges) between cases, resulting in swarms of connections that are difficult to interpret, an instance of the ‘hairball’ problem that plagues applications of networks to social and biological systems [43, 44]. In contrast, phylogenetic clusters are generally more interpretable as trees that are shaped in part by the underlying transmission history.

Both genetic distance and phylogenetic clustering methods require users to select one or more threshold parameters. In the absence of generic data-driven methods to select optimal thresholds, many users have resorted to default settings and/or emerging conventions in the literature [42, 45]. However, appropriate thresholds can vary substantially among databases and populations, due to differences in prevalence of infection, extent of sampling, and heterogeneity in rates of transmission and diagnosis [33, 46, 47]. In previous work [48], we developed a statistical framework to select the optimal threshold for distance-based clustering. This optimum is based on one’s ability to predict the distribution of the next cases of infection among existing clusters. Here, we extend this framework to phylogenetic clustering. This is not a trivial task because we must accommodate the effect of new data on the shape of the phylogenetic tree, such that new cases may retroactively change previous clusters. We adapt a maximum likelihood method (*pplacer* [49]) to graft new sequences onto a pre-existing phylogeny. Next, we fit predictive models of cluster growth based on the placement of new cases on the tree. To optimize the phylogenetic clustering method to a given data set, we evaluate these models for a range of branch-length and bootstrap support thresholds. We assess the performance of our method on HIV-1 sequence data sets from two regions of the United States (Tennessee [50] and Washington state [51]), the northern region of Alberta in Canada [52], and Beijing, China [53].

## METHODS

### Data collection

This study was performed on alignments of anonymized HIV-1 *pol* sequences, where each sequence uniquely represented a host individual. Aligned sequence data were obtained from different locations: Seattle, Washington, USA and the surrounding county (*n* = 6815) [51]; Middle Tennessee, USA (n = 2779) [50]; Beijing, China (n = 3964) [53]; and northern Alberta, Canada (n = 1054) [52]. The first three data sets were acquired with special permission from the Seattle/King County Public Health Agency [51, 54], the Vanderbuilt Comprehensive Care Clinic [15] and the Chinese CDC respectively. The fourth data set is publicly available in Genbank (accession numbers KU189996 – KU191050) [52] and we used a custom R script to extract collection dates and HIV-1 subtype classifications from sequence headers. Data sets were filtered to remove any sequences with over 5% ambiguous sites and sequences that were over 15% incomplete. We also manually examined and trimmed the sequence alignments for the Washington and Tennessee data sets due to a relatively high proportion of gaps and ambiguous base calls, removing a total of 52 and 163 nt from the overall alignment lengths, respectively. To make the results of our analysis as consistent as possible for the respective data contributors, we refrained from further modifications to the alignments.

All sequences were associated with years of sample collection as metadata. In addition, year of HIV diagnosis was available for all sequences in the Washington data set, and for a subset of sequences in the Tennessee data set (Table 1). For each data set, sequences were partitioned by date of sample collection, with sequences in the most recent year comprising the ‘incident’ subset, and the remaining sequences assigned to the ‘background’ subset. We partitioned the Tennessee data set into subsets by both year of sample collection and year of diagnosis, and repeated our analysis on both partitions. The composition of the final sequence data sets are summarized in Table 1.

**Table 1:**
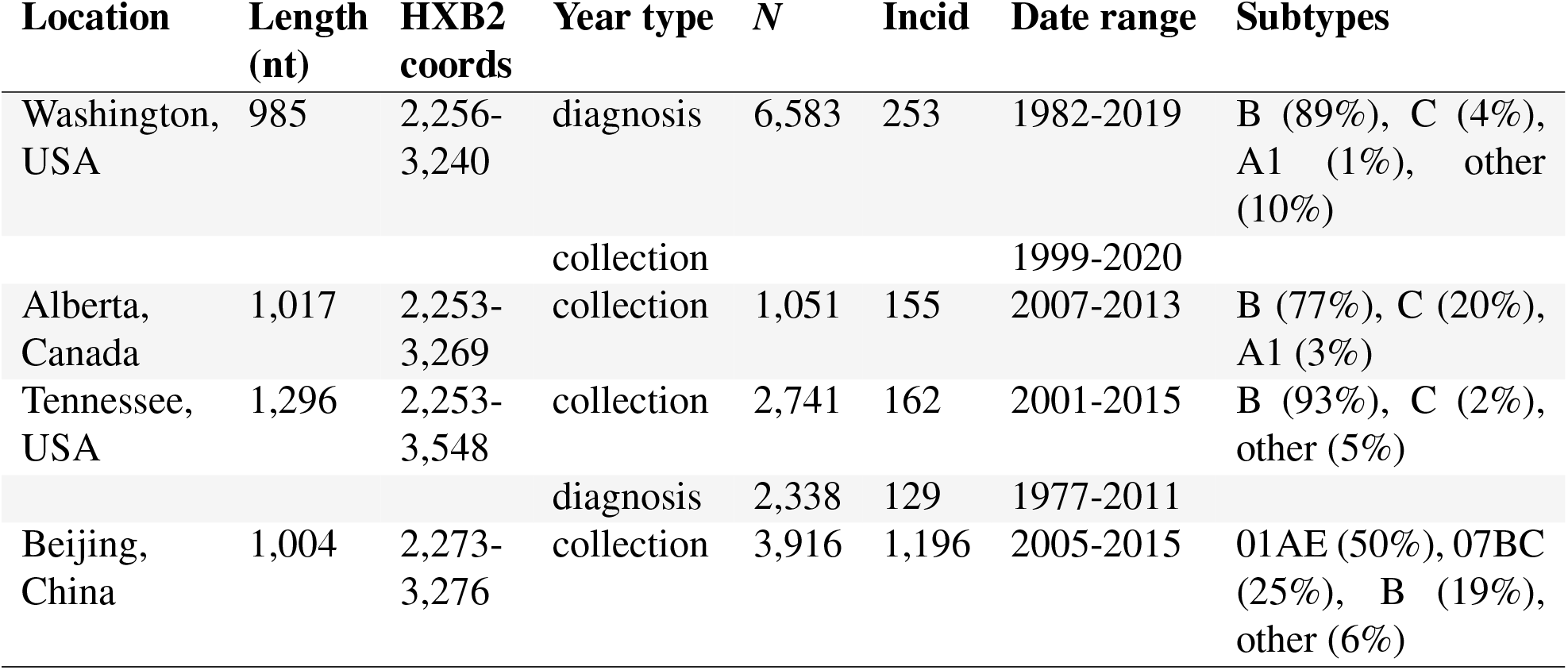
Summary of sequence data characteristics. Length is the median length of nucleotide (nt) sequences. HXB2 coords = reference nucleotide coordinates in the HXB2 genome (Genbank accession K03455). *N* = Total sample size, including both old and new sequences. Incid = number of sequences in ‘incident’ subset (most recent year). Subtype classifications were derived from the original data sources, when available, or generated *de novo* with SCUEAL [55].

| Location | Length (nt) | HXB2 coords | Year type | *N* | Incid | Date range | Subtypes |
| --- | --- | --- | --- | --- | --- | --- | --- |
| Washington, USA | 985 | 2,256-3,240 | diagnosis | 6,583 | 253 | 1982-2019 | B (89%), C (4%), A1 (1%), other (10%) |
|  |  |  | collection |  |  | 1999-2020 |  |
| Alberta, Canada | 1,017 | 2,253-3,269 | collection | 1,051 | 155 | 2007-2013 | B (77%), C (20%), A1 (3%) |
| Tennessee, USA | 1,296 | 2,253-3,548 | collection | 2,741 | 162 | 2001-2015 | B (93%), C (2%), other (5%) |
|  |  |  | diagnosis | 2,338 | 129 | 1977-2011 |  |
| Beijing, China | 1,004 | 2,273-3,276 | collection | 3,916 | 1,196 | 2005-2015 | 01AE (50%), 07BC (25%), B (19%), other (6%) |

### Phylogenetic analysis

Maximum likelihood phylogenies were constructed for each set of background sequences using IQ-TREE version 1.6.12 [56] with a generalized time-reversible (GTR) model of evolution, gamma-distributed rate variation among sites, and 1,000 parametric bootstraps. In addition, 100 random sub-samples were drawn without replacement from background sequences, each containing 80% of the full sample population. FastTree2 version 2.1.10 [57] was then used to construct a separate tree for each sub-sample, using default run parameters and 100 bootstraps. All trees were midpoint-rooted using the *phytools* [58] package in R. Polytomies were resolved arbitrarily using the *multi2di* function in the R package *ape* [59] — this was only necessary for sub-samples drawn from the Washington data set. Patristic distances (tip-to-tip branch lengths) were calculated using the R package *ape* function *cophenetic.phylo* and the pairwise distances were calculated using a Tamura-Nei [40] distance calculator (TN93, https://github.com/veg/tn93).

To update these phylogenies with incident sequences, we used the software package *pplacer* version 1.1 [49] to graft the sequences onto a tree by maximum likelihood. In many cases, the placement of a sequence varied among bootstrap replicates. The confidence levels among alternative placement queries (‘pqueries’) was quantified by likelihood weights. Each pquery is associated with a terminal branch between the incident sequence and the ancestral node *A* where it bisects the tree, as well as a ‘pendant’ branch between *A* and its descendant in the original tree (Figure 1). We used the *sing* subcommand in the *guppy* program within *pplacer* to generate trees with the different pqueries [60], ensuring that graft locations were only proposed on branches that existed in the original tree.

**Figure 1:**
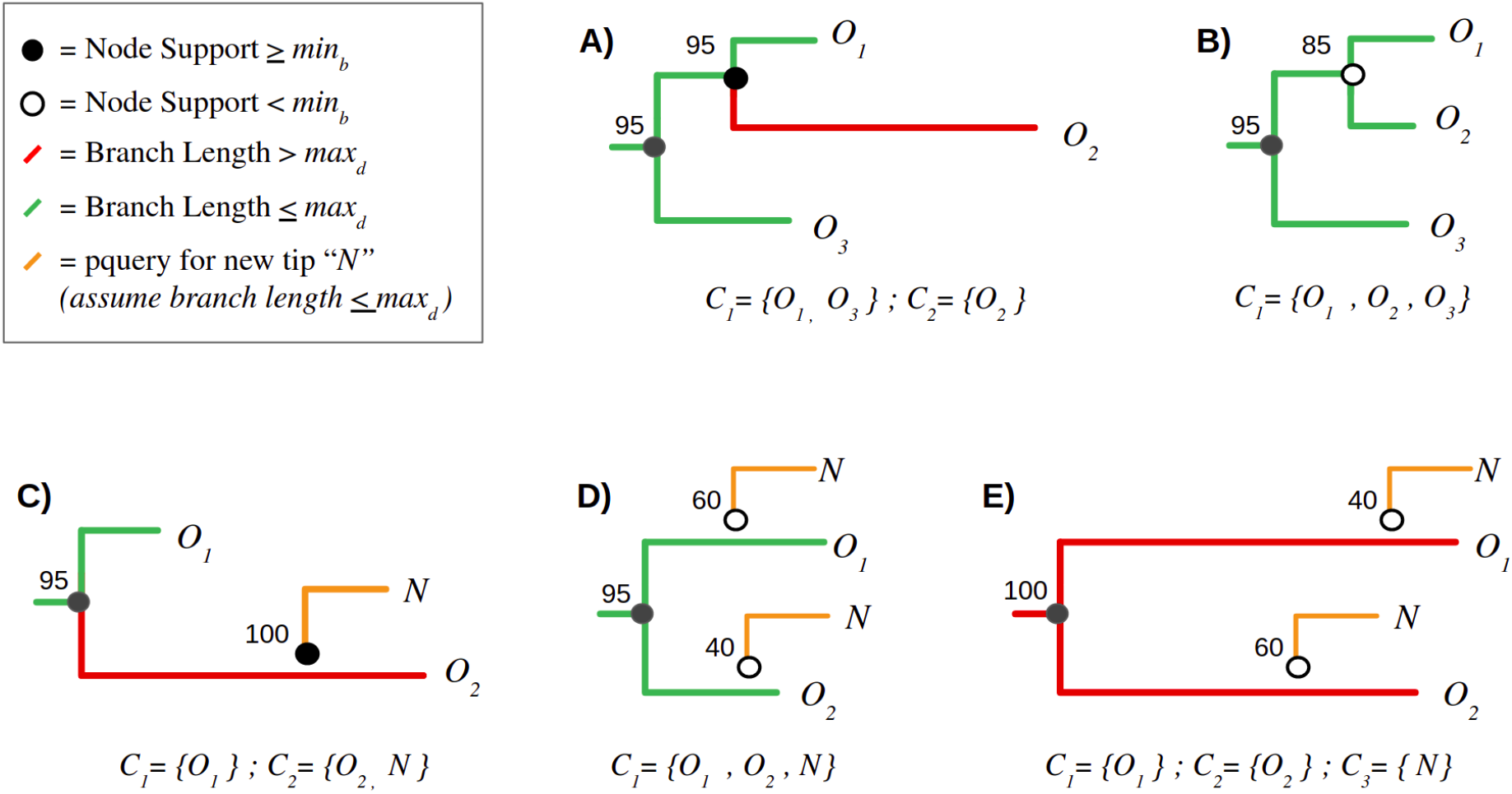
Examples of clustering definition and growth criteria. Subtree (A) represents a paraphyletic cluster (*C*_1_) of two background (old) sequences, *O*_1_ and *O*_3_, excluding a third sequence *O_2_* that is too distance from the root node of *C*_1_. Consequently, *O*_2_ becomes its own cluster of one, *C*_2_. Subtree (B) illustrates a monophyletic cluster where all background sequences meet this criterion. Subtrees (C)-(E) depict the addition of a new (incident) sequence *N* to an existing cluster. In (C), the new sequence is added with 100% confidence to a cluster of one background sequence, *O*_2_. Conversely, the placement of *N* in subtrees (D) and (E) is highly uncertain (with bootstrap supports 40% and 60%). For (D), *N* becomes incorporated into the same cluster irrespective of its placement, so the bootstrap values are irrelevant. In contrast, neither placement of *N* meets the clustering criteria due to the resulting branch lengths — as a result, *N* becomes a new cluster of one, *C*_3_.

### Cluster definition

Taking a phylogenetic tree of background sequences as input, we implemented the following set of rules in R to assign incident sequences to a set of clusters in the tree:

1. A cluster is a subset of branches and nodes within a binary tree rooted on ancestral node *A* and containing at least one terminal node.
2. By default, a single terminal node constitutes a cluster of one.
3. Each terminal node in a cluster must descend from the root node of the cluster through a path of branches whose lengths each fall below *d*_max_.
4. The ancestral node *A* must have a bootstrap confidence meeting or exceeding some threshold *b*_min_.
5. Any node *N* and its associated branch are only a member of the largest possible cluster containing *N*.

By these rules, a cluster can be as small as a single sequence, but can also be represented by either monophyletic or paraphyletic groups. The latter allows relatively divergent sequences to be separated from an otherwise closely related group, leaving the latter intact. Clusters with the same number of sequences can also vary by how many branches and internal nodes are included because of unresolved nodes (polytomies), although we do not utilize these quantities in our methods. We only count terminal nodes to compute the size of a cluster.

We generated clusters at 41 different branch length thresholds, ranging from *d*_max_ = 0 to 0.04 in steps of 0.001. Clusters were also generated under relaxed (*b*_min_ = 0) and strict (*b*_min_ = 0.95) bootstrap thresholds. Since our results were relatively insensitive to varying *b*_min_, we limited our analyses to these two settings. For the purpose of clustering, the root node of the tree was assigned a bootstrap support of 0, such that the entire tree could only become subsumed into a single giant cluster at the lowest threshold *b*_min_ = 0.

For comparison, we implemented a similar set of clustering criteria in the program Cluster Picker [27], which is widely used for the phylogenetic clustering analysis of HIV-1 sequences. This required two modifications to the above rules. First, a cluster selected by Cluster Picker must be a monophyletic group (containing all descendants) rooted on an ancestral node with a bootstrap support greater than or equal to a threshold *b*_min_. Second, all patristic distances in the cluster cannot exceed the threshold *d*_max_. We evaluated Cluster Picker at a broader range of distance thresholds, from *d*_max_ = 0 to 0.08 in steps of 0.002, and with *b*_min_ = 0.95. To distinguish our method from Cluster Picker, we will herein refer to the former as ‘paraphyletic clustering’.

### Simulation

We used *pplacer* to simulate the growth of clusters by the addition of incident sequences onto a fixed tree. This grafting creates new internal nodes and terminal branches and nodes. Next, we re-ran our paraphyletic clustering algorithm on the updated tree. To prevent any case where two clusters merge into one, only terminal or pendant branches descending from a newly created internal node were evaluated for clustering. It is possible for a single sequence to generate multiple proposed locations with varying confidence levels because *pplacer* employs bootstrap resampling. To avoid ambiguous placements, the internal node created by grafting a new sequence must have a bootstrap support that exceeds the threshold *b*_min_. If multiple placements with low support exist within the same cluster, we sum the support values to determine whether to assign the sequence to the cluster. We provide several examples of cluster growth in Figure 1 to clarify these definitions.

We also use *pplacer* to measure growth for the monophyletic clustering method in Cluster Picker. Unlike our paraphyletic clustering method, the addition of new tips in Cluster Picker has the potential to separate an existing cluster by introducing a new terminal branch with a length that exceeds *d*_max_. To prevent this kind of retroactive disruption to the composition of existing clusters, we only considered placements that would not result in a new distance over *d*_max_ for growth.

### Predictive performance analysis

Following our previous work [48], we used the predictive value of a Poisson regression model to optimize clustering parameters. In other words, we modelled the addition of incident cases to clusters of background sequences as a count outcome, with a cluster-specific rate *λ_i_* determined by the original composition of the i-th cluster. We propose that this approach is the most consistent with the implicit objectives of public health applications of clustering. We evaluated two Poisson models incorporating different sets of predictor variables. In the first model, cluster growth is predicted solely by the total number of background sequences in the cluster, also known as cluster size. This assumes that every individual background sequence has the same probability of being the closest relative to the next incident sequence. Thus, larger clusters tend to accumulate more new cases simply because they are large.

A second, more complex model incorporates an additional term corresponding to the mean ‘age’ or recency of sequences in the cluster relative to the current time. Age may be calculated from either dates of sample collection or diagnosis. Written more generally, we let *g*(*c*) = {0,1,2,…} represent the growth of cluster *c*:

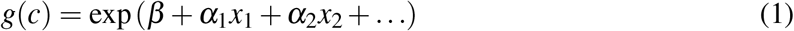

where *x_ι_* represents the *i*-th predictor variable, such as cluster size. Thus, this linear model can be modified to accommodate any number of predictor variables. The unit of observation for this model is a cluster of background sequences. For any model, we used the *glm* function in R to estimate the coefficients *α_i_* and intercept *β*.

We used the Akaike information criterion (AIC) [61] to compare the fit of alternative models on the same data. AIC increases both with model prediction error as well as the number of model parameters. To restate our central postulate, the optimal clustering threshold(s) define a partition of the data into clusters that maximizes the difference between the AIC of the null model, g_0_(*c*) = *β* + *α*_1_*x*_1_, which uses only cluster size (*x*_1_); and the AIC of an alternative model *g*_1_(*c*) = *β* + *α*_1_*x*_1_ + *α*_2_*x*_2_ that incorporates both cluster size and mean cluster age (*x*_2_). We measure this difference by Δ*AIC* = *AIC*(*g*_1_) – *AlC*(*g_0_*) and determine which combination of clustering thresholds *d*_max_ and *b*_min_ minimizes this quantity, *i.e*., attains the most negative value. At these optimized thresholds, the selection of a more informed model is the least ambiguous. We expect Δ*AIC* to approach zero at the most relaxed thresholds, where all background sequences are placed into a single giant cluster [48], such that it is not possible to differentiate between model outcomes. Conversely, at the strictest thresholds every background sequence is assigned to its own cluster of one. The predictive value of any characteristic such as age is diminished by extremely small sample size of each cluster, so we also expect Δ*AIC* to approach zero in this scenario.

In addition to the AIC-based approach, we used the R package pROC [62] to generate receiver operator characteristic (ROC) curves for varying cluster size and age thresholds. An ROC curve plots the true positive rate (TPR, sensitivity) and true negative rate (TNR, specificity) of a binary classifier when varying a single tuning parameter. Since our model predicts count-valued outcomes, we dichotomize the results to predict whether a cluster will grow by one or more cases *g*(*c*) > 0 or not *g*(*c*) = 0. For example, a true positive corresponds to a cluster that was both predicted and observed to accumulate one or more cases. Furthermore, we calculated the area under the curve (AUC) to provide a more conventional measure of model performance, where an AUC of 1 indicates perfect prediction.

## RESULTS

### Phylogenetic analysis

We reconstructed maximum likelihood phylogenies for alignments of HIV-1 *pol* sequences from each of four different sites (Washington and Tennessee, USA; Alberta, Canada; and Beijing, China). These alignments excluded all new (incident) sequences that were collected in the most recent year for each site. Figure 2 summarizes the distribution of branch lengths in the resulting trees. Although some of the data sets comprised multiple HIV-1 subtypes, this has a negligible effect on the internal branch length distributions because only a relatively small number of branches separate subtypes. Overall the distributions of internal and terminal branch lengths were broadly similar among locations. Internal branch lengths tended to be substantially shorter than terminal branch lengths (Figure 2), which is typical for HIV-1 among-host phylogenies. In addition, branches created by grafting new (incident) sequences onto the existing trees were significantly longer than terminal branches in the original trees (Wilcoxon test, *P* < 10^−10^; Figure 2). This trend may be driven by constraints imposed on the original tree when placing new sequences by maximum likelihood [49].

**Figure 2:**
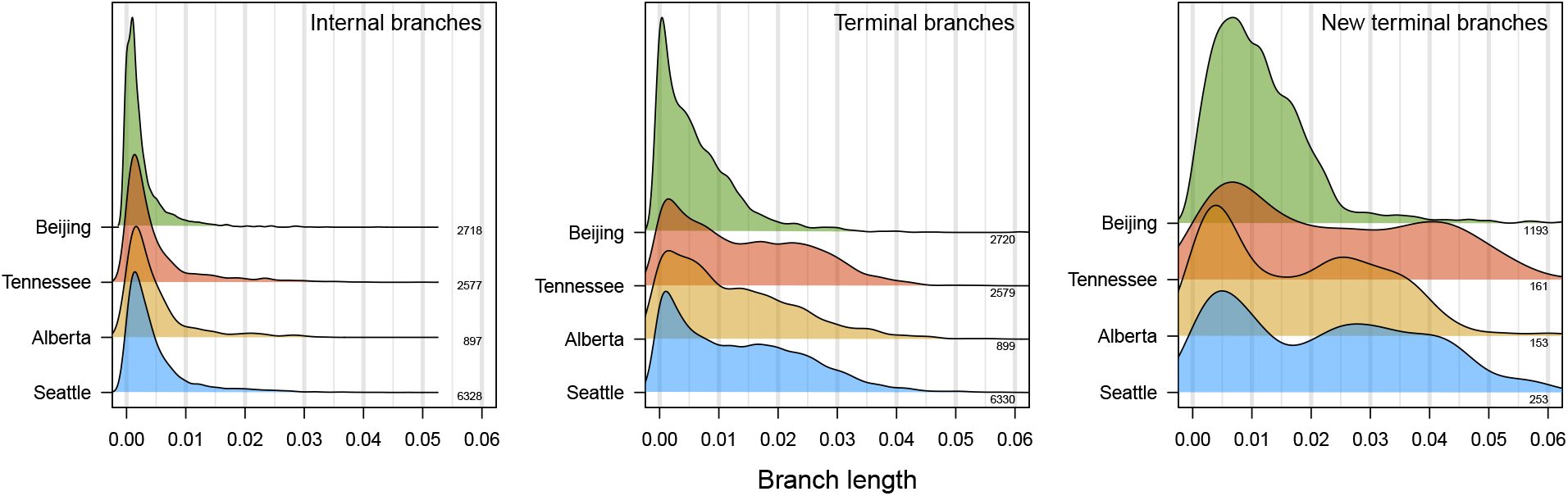
Distributions of branch lengths in HIV-1 *pol* phylogenies. Each density summarizes the distribution of branch lengths (measured in units of expected nucleotide substitutions per site) for different locations and types of branches, as indicated in the upper right corner of each plot. We used Gaussian kernel densities with default bandwidths adjusted by factors 1.5, 0.75 and 0.75, respectively. Densities are labeled on the right with the corresponding number of branches. Internal and terminal branches are derived from the phylogeny reconstructed from background sequences. New terminal branches refer to additional branches to incident (new) sequences as placed onto the phylogeny by maximum likelihood (pplacer).

The overall mean patristic distances were 0.102 (Washington), 0.115 (Alberta), 0.086 (Tennessee) and 0.159 (Beijing); the mean Tamura-Nei (TN93) distances ranged from 0.066 to 0.089, and were highly correlated with the patristic distances (Pearson’s *r*^2^ = 0.995). Differences in mean patristic distances among locations were driven in part by the presence of multiple major HIV-1 subtypes in the Alberta and Beijing phylogenies as distinct groups. For instance, the mean patristic distances within subtypes in the Beijing data set were 0.0630 (CRF01_AE), 0.0564 (CRF15) and 0.0564 (B). For Alberta, the mean patristic distances within subtypes were 0.067 (A1), 0.078 (B) and 0.074 (C). No cases of super-infection were reported in studies associated with these data sets, *e.g*., [50, 51, 53, 63].

### Predicting cluster growth

For each phylogeny of background sequences (excluding the most recent year), we extracted clusters under varying branch lengths (*d*_max_ from 0 to 0.04 in increments of 0.001 expected nucleotide substitutions per site) and bootstrap support (0% and 95%) thresholds. We measured cluster growth by the placement of new HIV-1 sequences (from the most recent year) onto the respective tree by maximum likelihood. As implied by the distribution of new terminal branch lengths (Figure 2), relaxing the branch length threshold (*d*_max_) resulted in higher rates of cluster growth. At the highest *d*_max_ evaluated in this study (0.04), 81% (Washington), 94% (Alberta), 77% (Tennessee) and 98% (Beijing) of all new sequences were grafted into existing clusters (Supplementary Figure S1). However, this high threshold also tended to collapse the background sequences into a single giant cluster, including sequences from different HIV-1 subtypes. Thus, at *d*_max_ exceeding 0.04, phylogenetic clusters are no more epidemiologically informative than the classification of the sequences into HIV-1 subtypes [64].

We modelled cluster growth as a Poisson-distributed outcome. Specifically, we evaluated different log-linked models using (1) the numbers of sequences per cluster, *i.e*., cluster size, or; (2) both cluster size and the mean times associated with sequences in clusters as predictor variables. We use the simplest case (cluster size only) as our null model. Times were based on either dates of sample collection or HIV diagnosis for the respective individuals. For instance, we expect a cluster comprising more recent infections to be more likely to gain new sequences. Fitting these models to a given data set provided two values of the Akaike information criterion (AIC), which measures the fit of the model penalized by the number of parameters. At a given set of thresholds defining clusters, the difference in AIC between these models quantifies the information gain by the addition of mean cluster times. At the most relaxed threshold, all sequences belong to a single cluster, and the addition of predictor variables has no effect on model fit. Conversely, at the strictest thresholds every background sequence is a cluster of one, such that the distribution of new sequences is essentially random with respect to individual characteristics. Thus, the impact of predictor variables on model information is contingent on how we partition the sample population.

The optimal clustering thresholds should resolve the bias-variance tradeoff between overfitting small clusters and underfitting large clusters. Put another way, the optimal thresholds minimize the information loss associated with the addition of one or more predictor variables, relative to the null model [48]. We quantify this information gain by computing the difference in AIC (ΔAIC) between the two models. Figure 3 illustrates the profiles of ΔAIC for the different data sets with respect to varying branch length (distance) thresholds with a fixed 95% bootstrap support threshold. In all four cases, ΔAIC was the most negative at intermediate distance thresholds, which varied slightly among locations: 0.007 (Alberta), 0.008 (Beijing), 0.012 (Tennessee), and 0.013 (Washington). We note that these optimal distance thresholds tend to be shorter than the thresholds often used in the literature (*e.g*., 0.045) after adjusting for the use of branch lengths (this study) versus tip-to-tip distances (e.g., Cluster Picker). The proportion of new sequences mapped to clusters at these optimal *d*_max_ thresholds were: 34.6% (Alberta), 40.2% (Beijing), 35.4% (Tennessee), and 39.1% (Washington; Supplementary Figure S1).

**Figure 3:**
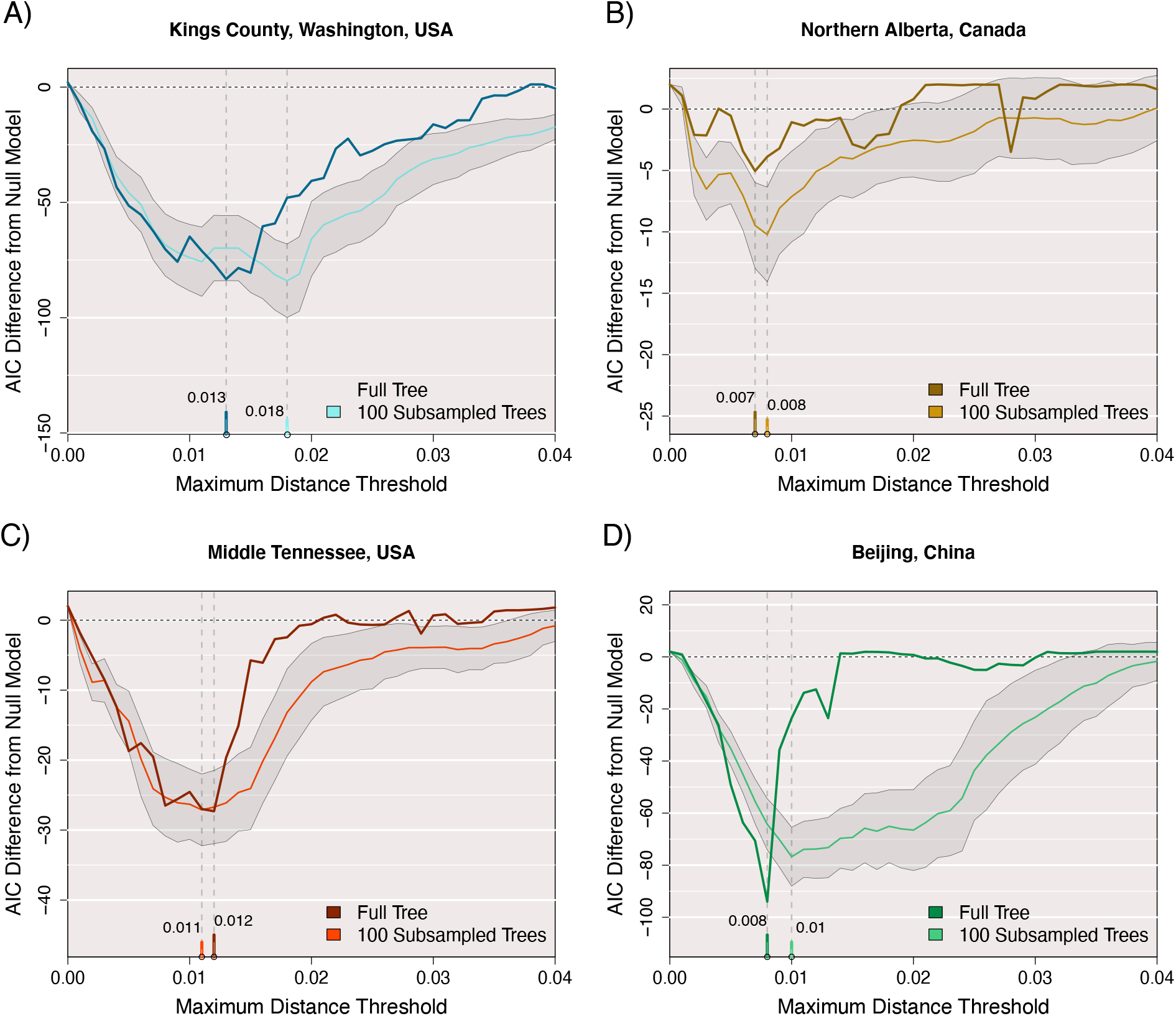
Difference in AIC between Poisson-linked models of cluster growth for all four full data sets. Clusters and growth are defined at 41 different maximum distance thresholds from 0 to 0.04 with a minimum bootstrap support requirement of 95% for ancestral nodes. The AIC of a null model where size predicts growth is subtracted from the AIC of a proposed model where size and mean time predict growth. The darker colour in each plot corresponds to these AIC results for a maxmimum likelihood tree built from the full set of old sequences, while the lighter colour represents the mean AIC difference obtained by this threshold for 100 approximate likelihood trees built on 80% subsamples of the old sequences without replacement. The shaded area represents 1 standard deviation from the mean AIC difference for subsamples at this threshold. Date of sequence collection was used to measure time for all data sets.

For a secondary measure of performance, the area under the curve (AUC) was obtained for receiver-operator characteristic (ROC) curves associated with the two Poisson regression models. In this case, cluster growth (using the paraphyletic method with *b*_min_ = 0.95) was reduced to a binary outcome. Put another way, we evaluated our ability to predict which clusters would grow by the addition of one or more new cases. At their respective optimal values of *d*_max_, we obtained AUC values of 0.87 (Seattle), 0.62 (Alberta), 0.76 (Tennessee) and 0.67 (Beijing) for the Poisson model using both cluster size and recency as predictor variables. These AUC values tended to be higher than the values obtained under the null model using only cluster size (Supplementary Figure S2). The exception was the Alberta data set, where similar AUC values were obtained from either model. In fact, the null model yielded significantly higher AUC values (paired Wilcoxon signed rank test, *P* = 0.01), and the AUCs were almost identical at the ΔAIC-selected *d*_max_ (0.629 null against 0.617 alternative). In contrast to our results for ΔAIC (Figure 3), the AUC profiles did not consistently select for an optimal *d*_max_ threshold, *i.e*., a global maximum. These results imply that AUC is not a reliable statistic for calibrating phylogenetic clustering methods on the basis of prospective cluster growth.

### Sensitivity analysis

To evaluate the sensitivity of these results to sample size and random variation, we sampled 80% of the background sequences at random without replacement to generate 100 replicates, and repeated our analysis for each sample. The distributions of the resulting optimal distance thresholds are summarized in Figure 3 and Supplementary Figure S3. The median opti-mal distance threshold for sub-samples of the Washington data was substantially greater (0.018) than the optimum for the full tree (0.013). This implied that the models required a more relaxed distance threshold to capture similar patterns of cluster growth with reduced sample sizes. However, subsetting the other data sets did not substantially change the optimal distance thresholds. Sub-sampling the Beijing data set resulted in an unusually broad ΔAIC profile, relative to the full data profile, and the distribution of optima among samples included a distinct ‘shoulder’ around a distance threshold of 0.02 (Supplementary Figure S3). Examining the cumulative plots of new sequences in clusters with increasing *d*_max_ (Supplementary Figure S1), we note that the optimal thresholds tended to coincide with the ‘elbow’ of the respective curves except for Beijing. This implies that the broader distribution of ΔAIC values associated with this data set may be driven by the atypically high number of incident sequences.

### Effect of bootstrap support

For all data sets, we generated alternative sets of AIC loss results with the minimum bootstrap threshold (*b*_min_) reduced to 0. The original threshold of *b*_min_ = 95% prevented many mid-sized clusters from forming. Specifically, 49% to 74% of internal nodes in each complete tree failed this threshold. This requirement also had a dramatic effect on the growth of singleton and small clusters, as 88% – 93% of the new sequence placements had *b* < 95%, often resulting in clusters growing only through multiple placements. Overall, relaxing *b*_min_ tended to reduce the loss of information, as reflected by lower values of ΔAIC (Figure 4). This outcome was the most apparent in the Washington data set, which had the largest proportion of internal nodes above *b*_min_ = 95%. Relaxing *b*_min_ also tended to induce a shift in the optimal *d*_max_ as defined by minimizing ΔAIC, although this appeared to be a stochastic outcome. In contrast, relaxing *b*_min_ for the Beijing data set resulted in higher values of ΔAIC in the neighbourhood of the optimal *d*_max_ threshold. Again we attribute this difference to the atypically high number of incident cases for this data set. In sum, our results imply that relaxing the threshold *b*_min_ tends to confer a prediction advantage by incorporating a greater number of clusters into the analysis, even though many of those clusters would not be consistently reproducible with new data.

**Figure 4:**
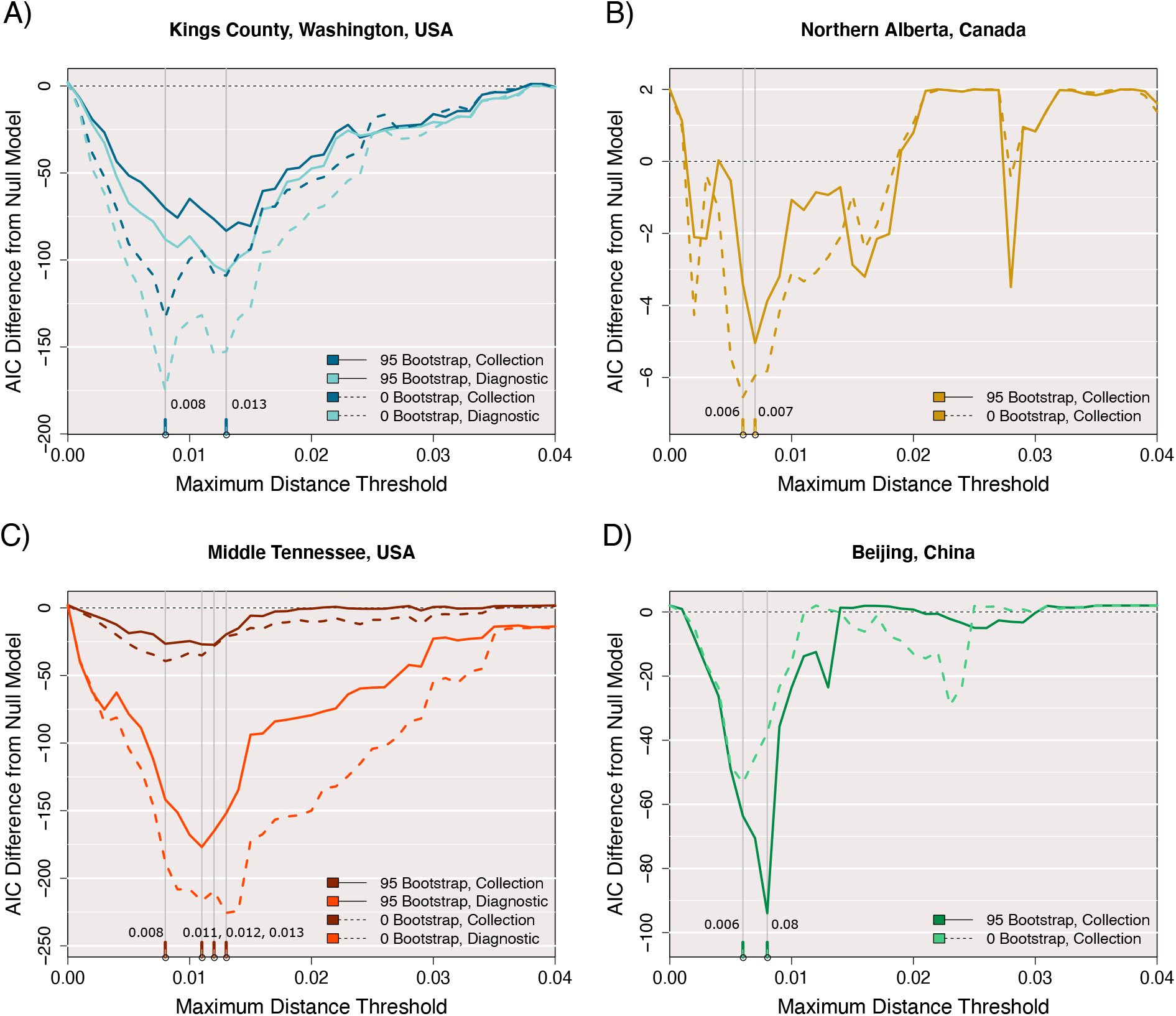
The AIC difference between two Poisson-linked models of cluster growth for all four full data sets, with clusters and growth defined at 41 different maximum distance thresholds from 0 to 0.04. The AIC of a null model where size predicts growth is subtracted from the AIC of a proposed model where size and time predict growth. For the data sets where either sequence collection date or associated patient diagnostic date could define time, both AIC difference results are shown by separate colours. Solid lines represent AIC differences obtained using an additional bootstrap threshold of 0.95 to define clusters and growth, while dashed lines were obtained without this requirement.

### Diagnosis versus collection dates

For the Washington and Tennessee data sets, we generated an additional set of ΔAIC profiles cluster ages computed from the available dates of HIV diagnosis instead of sample collection dates. Given that dates of HIV diagnosis tend to be closer to the actual dates of infection, we expect these metadata to be more useful for predicting the distribution of new cases. Indeed, using the diagnosis dates resulted in substantially lower ΔAIC values in either data set, irrespective of bootstrap thresholding (Figure 4). The optimal *d*_max_ as determined by ΔAIC was invariant to using either set of dates for the Washington data set. On the other hand, we obtained slightly different optima for Tennessee, most likely because diagnostic dates were only available for a subset of incident cases in this data set (80%, Table 1).

### Monophyletic clustering

Finally, we generated another set of ΔAIC profiles using clusters that were constrained to be monophyletic, *i.e*., comprising all descendants of the internal node. This second clustering method is more similar to the phylogenetic clustering methods used in the molecular epidemiology literature. For example, Cluster Picker [19] defines clusters as monophyletic clades with a default bootstrap requirement of ≥ 95% and a maximum patristic distance between all sequences within the subtree. Supplementary Figure S4 displays the ΔAIC profiles for all four data sets across 41 different patristic distance thresholds and *b*_min_ = 0.95. One notable feature of these profiles is that the ΔAIC values do not consistently converge to zero at a distance threshold of zero. The ΔAIC profiles for the Alberta and Tennessee data sets were visibly less responsive to variation in distance thresholds, making it difficult to estimate optimal thresholds.

Table 2 summarizes several clustering statistics for paraphyletic and monophyletic clustering, obtained at their respective optimal *d*_max_ thresholds. As noted in the preceding section, removing the bootstrap threshold substantially increases the number of acceptable clusters, while the ΔAIC optima consistently shifted to slightly lower values of *d*_max_. In addition, relaxing *b*_min_ tends to increase the total number of incident sequences that are connected to clusters. For the Beijing data set, however, the number of incident sequences is reduced by setting *b*_min_ = 0; we attribute this to the exclusion of incident cases in paraphyletic clusters by the stricter *d*_max_ threshold selected by ΔAIC. Under monophyletic clustering, the distribution of cluster sizes was more constrained, as it becomes increasingly likely that larger portions of the tree incorporate one or more branch lengths that causes the maximum in-cluster patristic distance to exceed the threshold.

**Table 2:** Cluster statistics under paraphyletic and monophyletic clustering. ‘No bootstrap’ corresponds to paraphyetic clustering with *b*_min_ = 0; otherwise this threshold defaults to 95%. Optimal *d*_max_ is the pairwise distance threshold selected by minimizing ΔAIC, in units of expected number of nucleotide substitutions. ‘Number of clusters’ only counts clusters with two or more background sequences, *i.e*., this number does not include singletons. Total growth is the number of incident (new) sequences connected to clusters of background sequences. Growing clusters is the number of clusters to which incident sequences attach.

| Location | Optimal<br>$d_{\max}$ | Number of<br>clusters > 1 | Mean<br>cluster size | Largest<br>cluster size | Total<br>growth | Growing<br>clusters |
| --- | --- | --- | --- | --- | --- | --- |
| Washington, USA | 0.013 | 74 | 1.39 | 1657 | 68 | 30 |
| *No bootstrap* | 0.008 | 378 | 2.04 | 1560 | 73 | 36 |
| *Monophyletic* | 0.016 | 639 | 1.20 | 16 | 92 | 96 |
| Alberta, Canada | 0.007 | 104 | 1.35 | 58 | 34 | 27 |
| *No bootstrap* | 0.006 | 228 | 1.75 | 28 | 45 | 28 |
| *Monophyletic* | 0.002 | 26 | 1.04 | 4 | 52 | 64 |
| Tennessee, USA | 0.012 | 1 | 1.35 | 358 | 41 | 28 |
| *No bootstrap* | 0.008 | 621 | 1.91 | 78 | 41 | 35 |
| *Monophyletic* | 0.020 | 360 | 1.29 | 11 | 63 | 53 |
| Beijing, China | 0.008 | 135 | 1.82 | 533 | 373 | 119 |
| *No bootstrap* | 0.006 | 409 | 2.49 | 381 | 305 | 127 |
| *Monophyletic* | 0.032 | 490 | 2.13 | 46 | 613 | 275 |

## DISCUSSION

Genetic clustering results can vary substantially with different methods and clustering thresholds [45]. There are no universal guidelines for configuring a phylogenetic clustering method for applications in public health and molecular epidemiology. Here, we have described a means of calibrating phylogenetic clustering to a specific population database, based on our ability to predict where the next cases will appear. Relaxing clustering thresholds tend to yield larger clusters [19, 37, 38]. Several studies [31, 33, 35, 65] have noted that larger clusters tend to have more information for predicting the location of the next cases. However, this favours relaxing the clustering thresholds until all sequences merge to a single giant cluster, and the appearance of the next cases in that cluster is a trivial outcome [48]. An analogous information-bias tradeoff has been studied extensively in spatial statistics, where it is known as the modifiable area unit problem [66, 67]. Thus, we have adapted a strategy that was proposed to study the distribution of mortality rates for varying levels of administrative districts in Tokyo [68] (e.g., ward, town, village). This method compares the AICs of two Poisson regression models for varying numbers of clusters. Although AICs for one model cannot be directly compared for different clusterings (since the data are being changed), the clustering that maximizes the difference in AICs between models (ΔAIC) minimizes information loss with the addition of model parameters [69, 70].

In our results, ΔAIC consistently approached zero at the extremes of the distance threshold (*d*_max_), indicating that the addition of sample collection or diagnosis dates had no impact on our ability to predict the next cases under these extreme clustering settings. The intermediate *d*_max_ values that minimized the ΔAIC varied among data sets and subsamples of the same data. Selected thresholds generally fell within the range of *d*_max_ settings used frequently in the literature — adjusting for node-to-tip versus tip-to-tip measures, this range roughly spans 0.0075 to 0.0225. For example, the original study associated with the Seattle data set [51] had employed monophyletic clustering (Cluster Picker) with *b*_min_ = 0.95 and *d*_max_ = 0.0075. In contrast, our analysis favoured a more relaxed threshold for monophyletic clusters in this data set (*d*_max_ = 0.016; Table 2). However, ΔAIC was also relatively invariant to changes in *d*_max_ under these conditions (Supplementary Figure S4).

In contrast, the effect of *b*_min_ on phylogenetic clustering has not received as much attention as *d*_max_, possibly due to the popularity of (non-phylogenetic) distance-based clustering methods such as HIV-TRACE [7]. Novitsky and colleagues [71] recently reported that reducing the bootstrap threshold resulted in smaller and more numerous clusters. However, they evaluated a smaller range of bootstrap thresholds (*b*_min_ = 0.7 — 1.0) and employed no distance criterion for their analysis of HIV-1 subtype C sequences from a diverse number of locations, including South Africa, Botswana and India. In our analysis, we found that ΔAIC was maximized when phylogenetic clusters were generated under no bootstrap requirement, *i.e*., *b*_min_ = 0, which tended to yield greater numbers of small clusters with no substantial impact on selecting *d*_max_ (Table 2). For comparison, the study originally associated with the northern Alberta data set [52] defined clusters as monophyletic clades with *b*_min_ = 0.95. Additionally, they applied a distance criterion to a time-scaled tree, such that lengths were measured in units of time, i.e., *d*_max_ =5–10 years. This was an interesting choice, because our analysis of monophyletic clustering on their data resulted in ΔAIC close to zero across a range of *d*_max_ measured as a genetic distance (expected number of substitutions per site; Supplementary Figure S4).

Rose and colleagues [35] recently also explored the selection of clustering thresholds for phylogenetic methods in application to HIV-1. Specifically, they evaluated the ability of Cluster Picker [27] to place known HIV-1 transmission pairs (i.e., epidemiologically linked heterosexual couples in the Rakai Cohort Community Study) into clusters under varying distance thresholds. They determined that *d*_max_ between 0.04 and 0.053 were the most effective for distinguishing between epidemiologically linked and unlinked pairs in their study population. We note that their application of clustering is markedly different from our study, which focuses on the distribution of incident cases at a population level. Hence, we make no attempt to infer direct transmission between individuals, which has significant ethical and legal implications [72], and our results are unlikely to be useful for that application. Studies that focus on direct transmission require fundamentally different sampling strategies as a cluster may represent indirect transmission through an unsampled intermediary [46, 73].

Retrospective studies of phylogenetic clusters of HIV-1 sequences are common [19, 51, 74]. Clusters are used as a proxy for variation in transmission rates that may be statistically associated with different risk factors, such as injection drug use. In this context, defining clusters is relatively straightforward. Tracking clusters prospectively over time, however, raises significant issues for phylogenetic clustering because a cluster can be broken into a set of smaller clusters by the addition of new cases. Distance clustering methods do not suffer from this problem because pairwise distances are invariant to the addition of new data [16, 39]. This distinction is similar to the difference between single-linkage and complete-linkage clustering [36]. To circumvent this limitation of phylogenetic clustering, we have modified the conventional definition (*e.g*., Cluster Picker [27], Tree Cluster [30]) by incorporating the concept of grafting new sequences onto a tree [49]. This allows clusters to exclude long branch lengths, making it more similar to parameteric methods for phylogenetic clustering that have recently been proposed [28, 29]. An interesting challenge with this redefinition of phylogenetic clusters, however, is that relatively long terminal branch lengths can disqualify tips from ever becoming assigned to clusters. Variation in terminal branch lengths can be driven by differences in sampling/diagnosis rates among groups [46, 47]. In addition, the discordance between rates of HIV-1 evolution within and among hosts [63] can contribute to terminal branch lengths in the virus phylogeny.

Our study has focused on the benefit of knowing the times since sample collection or diagnosis on our ability to predict the location of the next infections. Although other risk factors such as injection drug use or commercial sex work are often associated with cluster formation [22, 51, 74] and growth [16, 38, 39], sampling times are the most consistently available metadata for HIV-1 sequences. Since the Poisson regression model can hypothetically accommodate any number of predictor variables, we expect that incorporating additional metadata will drive a further decrease in ΔAIC and may shift the location of the optimal *d*_max_. Although ΔAIC may offer a framework for variable selection in the context of cluster optimization [48], the statistical justification for this is unclear and remains an area for further work. For example, we cannot directly compare AIC values obtained from different models at different clustering thresholds, because the underlying data have changed. Therefore, variable selection may not be as simple as determining which model minimizes the ΔAIC across thresholds. However, our results support the use of the AIC loss metric and prospective growth modeling to adjust phylogenetic clustering studies for the genetic composition of each study population. This method can also play an important role in extending these tools to other viruses, such as the ongoing SARS-CoV-2 pandemic [2, 3, 71, 75].

## ACKNOWLEDGEMENTS

We thank Dr. Susan Buskin and Richard Lechtenberg for their work at Public Health - Seattle & King County that made this study possible. This study was supported by a grant from the Canadian Institutes for Health Research (PJT-156178) and by an Administrative Supplement to a Center Core Grant from the Tennessee Center for AIDS Research (P30-AI110527).

## SUPPLEMENTARY FIGURES

**Figure S1:**
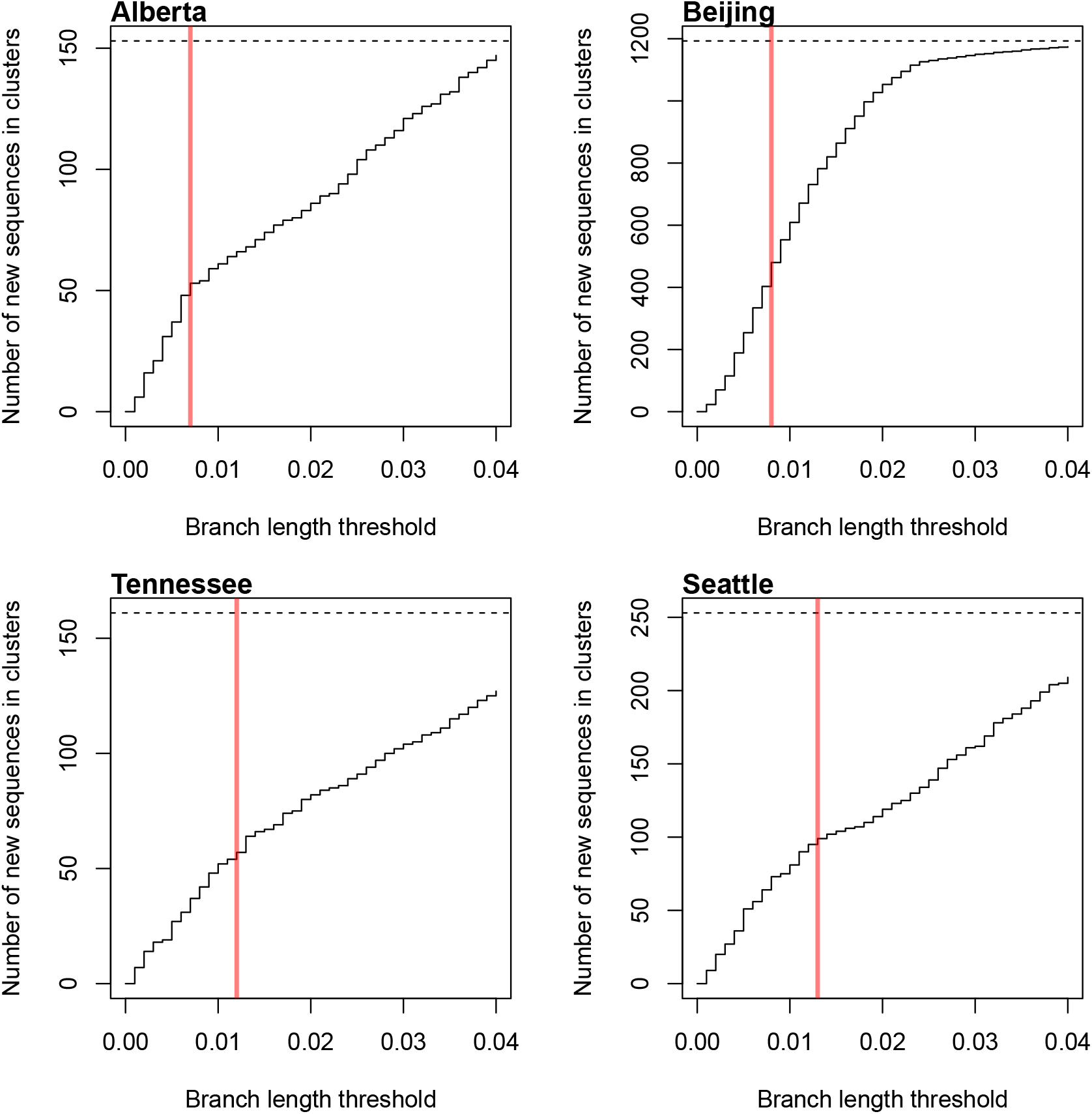
Step charts of the number of incident (new) HIV-1 sequences mapped to clusters in the original phylogeny, as a function of the distance threshold *d*_max_. For each data set, the total number of new sequences is indicated by a horizontal dashed line, and the optimal *d*_max_ is represented by a vertical solid red line.

**Figure S2:**
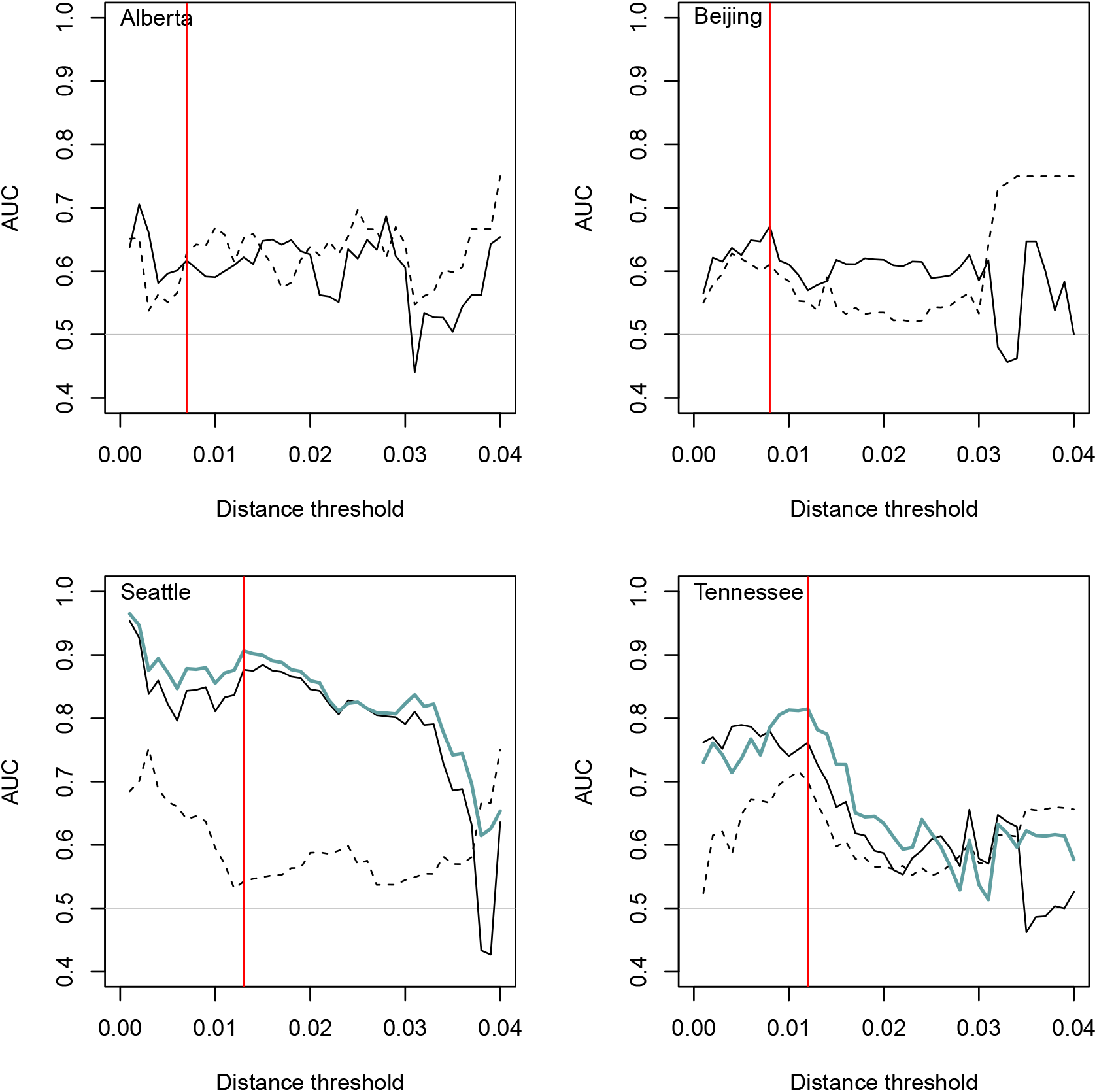
Summary of AUC (area under the curve) results. AUC values were calculated from the receiver-operator characteristic curves associated with predicting the attachment of one or more new cases to known clusters. Dashed black lines represent AUC values obtained under the null model (cluster size only). Solid black lines represent AUC values obtained under the alternative model (cluster size and age based on dates of sample collection). Solid thick blue lines represent AUC values obtained under the alternative model with dates of HIV diagnosis, when available. A faint horizontal line is drawn at AUC=0.5 where a model is no better than a random guess, and a red vertical line is drawn at the *d*_max_ threshold selected by ΔAIC.

**Figure S3:**
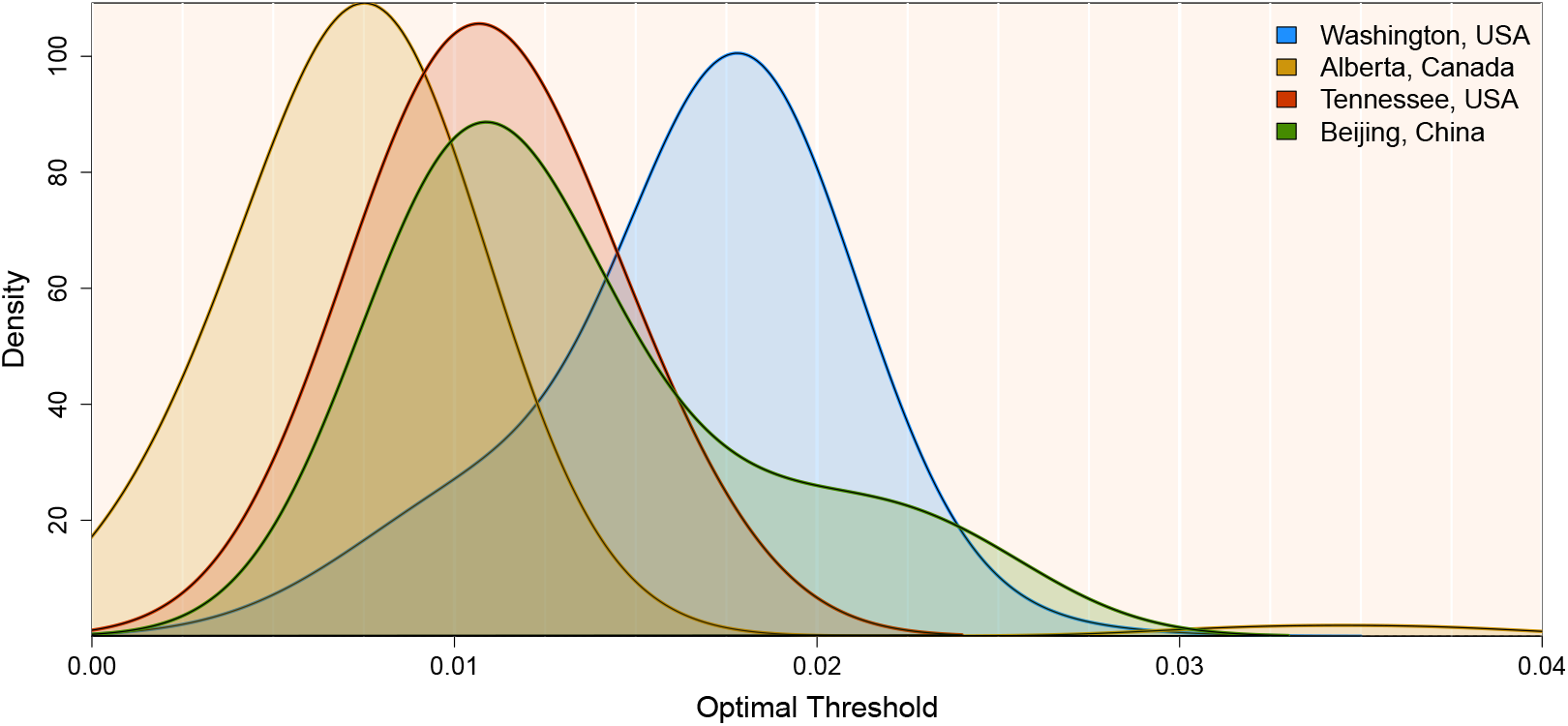
The density of optimal *d*_max_ values for all 4 data sets, based on approximate maximum likelihood trees built from 100 random 80% subsamples of each data set without replacement. A bandwidth of 0.003 was used for consistency across data sets. Optimal *d*_max_ values were defined as those which lead to clusters and growth results that yielded the greatest loss of AIC when comparing two Poisson-linked models of cluster growth: a null model where size predicts growth, and a proposed model where size and time predict cluster growth. For all trees, a minimum bootstrap requirement of 95% was used to define clusters.

**Figure S4:**
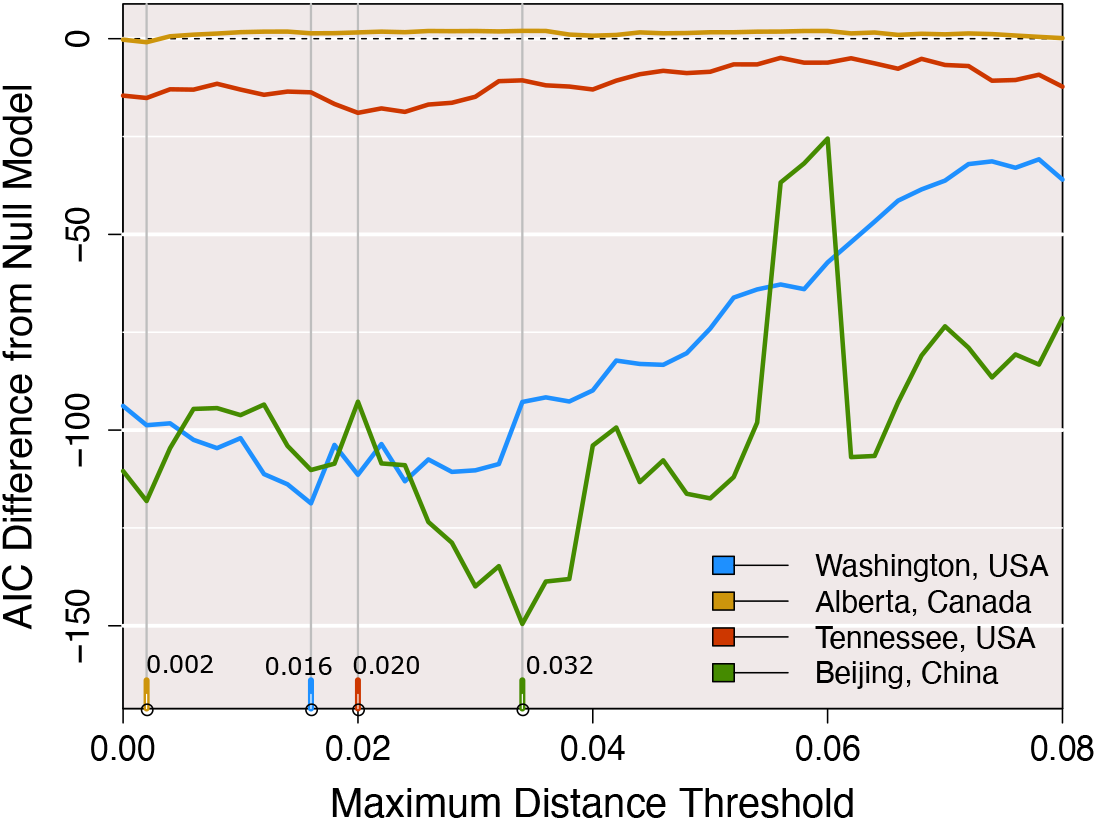
The AIC difference between two Poisson-linked models of cluster growth for all four full data sets, with monophyletic clusters and growth defined at 41 different maximum patristic distance thresholds from 0 to 0.04. The AIC of a null model where size predicts growth is subtracted from the AIC of a proposed model where size and time predict growth. Unlike previous figures, these loss values were obtained using an alternative method of clustering.

## REFERENCES

[1] Zhu N, Zhang D, Wang W, Li X, Yang B, Song J, et al. A novel coronavirus from patients with pneumonia in China, 2019. New England Journal of Medicine. 2020;382:727–733.

[2] Furuse Y, Sando E, Tsuchiya N, Miyahara R, Yasuda I, Ko YK, et al. Clusters of coronavirus disease in communities, Japan, January–April 2020. Emerging infectious diseases. 2020;26(9):2176.

[3] Pung R, Chiew CJ, Young BE, Chin S, Chen MI, Clapham HE, et al. Investigation of three clusters of COVID-19 in Singapore: implications for surveillance and response measures. The Lancet. 2020;395(10229):1039–1046.

[4] Pini A, Zomahoun D, Duraffour S, Derrough T, Charles M, Quick J, et al. Field investigation with real-time virus genetic characterisation support of a cluster of Ebola virus disease cases in Dubréka, Guinea, April to June 2015. Eurosurveillance. 2018;23(12):17–00140.

[5] Gire SK, Goba A, Andersen KG, Sealfon RS, Park DJ, Kanneh L, et al. Genomic surveillance elucidates Ebola virus origin and transmission during the 2014 outbreak. Science. 2014;345(6202):1369–1372.

[6] Poon AF, Gustafson R, Daly P, Zerr L, Demlow SE, Wong J, et al. Near real-time monitoring of HIV transmission hotspots from routine HIV genotyping: an implementation case study. The lancet HIV. 2016;3(5):e231–e238.

[7] Kosakovsky Pond SL, Weaver S, Leigh Brown AJ, Wertheim JO. HIV-TRACE (TRAns-mission Cluster Engine): a tool for large scale molecular epidemiology of HIV-1 and other rapidly evolving pathogens. Molecular biology and evolution. 2018;35(7):1812–1819.

[8] Volz EM, Le Vu S, Ratmann O, Tostevin A, Dunn D, Orkin C, et al. Molecular epidemiology of HIV-1 subtype B reveals heterogeneous transmission risk: implications for intervention and control. The Journal of infectious diseases. 2018;217(10):1522–1529.

[9] Han WM, Colby DJ, Khlaiphuengsin A, Apornpong T, Kerr SJ, Ubolyam S, et al. Large transmission cluster of acute hepatitis C identified among HIV-positive men who have sex with men in Bangkok, Thailand. Liver International. 2020;40(9):2104–2109.

[10] Zumla A, Hui DS, Perlman S. Middle East respiratory syndrome. The Lancet. 2015;386(9997):995–1007.

[11] Zhong N, Zheng B, Li Y, Poon L, Xie Z, Chan K, et al. Epidemiology and cause of severe acute respiratory syndrome (SARS) in Guangdong, People’s Republic of China, in February, 2003. The Lancet. 2003;362(9393):1353–1358.

[12] Drake JW, Holland JJ. Mutation rates among RNA viruses. Proceedings of the National Academy of Sciences. 1999;96(24):13910–13913.

[13] Moya A, Holmes EC, González-Candelas F. The population genetics and evolutionary epidemiology of RNA viruses. Nature Reviews Microbiology. 2004;2(4):279–288.

[14] Ypma RJ, van Ballegooijen WM, Wallinga J. Relating phylogenetic trees to transmission trees of infectious disease outbreaks. Genetics. 2013;195(3):1055–1062.

[15] Dennis AM, Hue S, Billock R, Levintow S, Sebastian J, Miller WC, et al. Human immunod-eficiency virus type 1 phylodynamics to detect and characterize active transmission clusters in North Carolina. The Journal of Infectious Diseases. 2020;221(8):1321–1330.

[16] Billock RM, Powers KA, Pasquale DK, Samoff E, Mobley VL, Miller WC, et al. Prediction of HIV transmission cluster growth with statewide surveillance data. Journal of acquired immune deficiency syndromes (1999). 2019;80(2):152.

[17] De Oliveira T, Kharsany AB, Gräf T, Cawood C, Khanyile D, Grobler A, et al. Transmission networks and risk of HIV infection in KwaZulu-Natal, South Africa: a community-wide phylogenetic study. The lancet HIV. 2017;4(1):e41–e50.

[18] Dalai SC, Junqueira DM, Wilkinson E, Mehra R, Kosakovsky Pond SL, Levy V, et al. Combining Phylogenetic and Network Approaches to Identify HIV-1 Transmission Links in San Mateo County, California. Frontiers in microbiology. 2018;9:2799.

[19] Ragonnet-Cronin M, Lycett SJ, Hodcroft EB, Hue S, Fearnhill E, Brown AE, et al. Transmission of non-B HIV subtypes in the United Kingdom is increasingly driven by large non-heterosexual transmission clusters. The Journal of infectious diseases. 2016;213(9):1410–1418.

[20] Kiwuwa-Muyingo S, Nazziwa J, Ssemwanga D, Ilmonen P, Njai H, Ndembi N, et al. HIV-1 transmission networks in high risk fishing communities on the shores of Lake Victoria in Uganda: A phylogenetic and epidemiological approach. PLoS One. 2017;12(10):e0185818.

[21] Charre C, Cotte L, Kramer R, Miailhes P, Godinot M, Koffi J, et al. Hepatitis C virus spread from HIV-positive to HIV-negative men who have sex with men. PLoS One. 2018;13(1):e0190340.

[22] Sivay MV, Palumbo PJ, Zhang Y, Cummings V, Guo X, Hamilton EL, et al. HIV drug resistance, phylogenetic analysis, and superinfection among men who have sex with men and transgender women in sub-Saharan Africa: HPTN 075. Clinical Infectious Diseases. 2020;73(1):50–59.

[23] Fogel JM, Sivay MV, Cummings V, Wilson EA, Hart S, Gamble T, et al. HIV drug resistance in a cohort of HIV-infected MSM in the United States. Aids. 2020;34(1):91–101.

[24] Grulich AE, Guy R, Amin J, Jin F, Selvey C, Holden J, et al. Population-level effectiveness of rapid, targeted, high-coverage roll-out of HIV pre-exposure prophylaxis in men who have sex with men: the EPIC-NSW prospective cohort study. The lancet HIV. 2018;5(11):e629–e637.

[25] Masyuko S, Mukui I, Njathi O, Kimani M, Oluoch P, Wamicwe J, et al. Pre-exposure prophylaxis rollout in a national public sector program: the Kenyan case study. Sexual health. 2018;15(6):578–586.

[26] Fauci AS, Redfield RR, Sigounas G, Weahkee MD, Giroir BP. Ending the HIV epidemic: a plan for the United States. Jama. 2019;321(9):844–845.

[27] Ragonnet-Cronin M, Hodcroft E, Hué S, Fearnhill E, Delpech V, Brown AJL, et al. Auto-mated analysis of phy-logenetic clusters. BMC bioinformatics. 2013;14(1):317.

[28] Barido-Sottani J, Vaughan TG, Stadler T. Detection of HIV transmission clusters from phylogenetic trees using a multi-state birth–death model. Journal of the Royal Society Interface. 2018;15(146):20180512.

[29] McCloskey RM, Poon AF. A model-based clustering method to detect infectious disease transmission outbreaks from sequence variation. PLoS computational biology. 2017;13(11):e1005868.

[30] Balaban M, Moshiri N, Mai U, Jia X, Mirarab S. TreeCluster: Clustering biological sequences using phylogenetic trees. PloS one. 2019;14(8):e0221068.

[31] Han AX, Parker E, Maurer-Stroh S, Russell CA. Inferring putative transmission clusters with Phydelity. Virus Evolution. 2019;5(2):vez039.

[32] Prosperi MC, Ciccozzi M, Fanti I, Saladini F, Pecorari M, Borghi V, et al. A novel methodology for large-scale phylogeny partition. Nature communications. 2011;2(1):1–10.

[33] Poon AF. Impacts and shortcomings of genetic clustering methods for infectious disease outbreaks. Virus evolution. 2016;2(2):vew031.

[34] Berry V, Gascuel O. On the interpretation of bootstrap trees: appropriate threshold of clade selection and induced gain. Molecular Biology and Evolution. 1996;13(7):999–1011.

[35] Rose R, Lamers SL, Dollar JJ, Grabowski MK, Hodcroft EB, Ragonnet-Cronin M, et al. Identifying transmission clusters with cluster picker and HIV-TRACE. AIDS research and human retroviruses. 2017;33(3):211–218.

[36] Bbosa N, Ssemwanga D, Kaleebu P. Choosing the right program for the identification of HIV-1 transmission networks from nucleotide sequences sampled from different populations. AIDS Research and Human Retroviruses. 2020;36(11):948–951.

[37] Erly SJ, Herbeck JT, Kerani RP, Reuer JR. Characterization of Molecular Cluster Detection and Evaluation of Cluster Investigation Criteria Using Machine Learning Methods and Statewide Surveillance Data in Washington State. Viruses. 2020;12(2):142.

[38] Oster AM, France AM, Panneer N, Ocfemia MCB, Campbell E, Dasgupta S, et al. Identifying clusters of recent and rapid HIV transmission through analysis of molecular surveillance data. Journal of acquired immune deficiency syndromes (1999). 2018;79(5):543.

[39] Wertheim JO, Murrell B, Mehta SR, Forgione LA, Kosakovsky Pond SL, Smith DM, et al. Growth of HIV-1 molecular transmission clusters in New York City. The Journal of infectious diseases. 2018;218(12):1943–1953.

[40] Tamura K, Nei M. Estimation of the number of nucleotide substitutions in the control region of mitochondrial DNA in humans and chimpanzees. Molecular biology and evolution. 1993;10(3):512–526.

[41] Poon AF, Joy JB, Woods CK, Shurgold S, Colley G, Brumme CJ, et al. The impact of clinical, demographic and risk factors on rates of HIV transmission: a population-based phylogenetic analysis in British Columbia, Canada. The Journal of infectious diseases. 2015;211(6):926–935.

[42] Hassan AS, Pybus OG, Sanders EJ, Albert J, Esbjörnsson J. Defining HIV-1 transmission clusters based on sequence data. AIDS (London, England). 2017;31(9):1211.

[43] Dianati N. Unwinding the hairball graph: pruning algorithms for weighted complex networks. Physical Review E. 2016;93(1):012304.

[44] Röttjers L, Faust K. From hairballs to hypotheses–biological insights from microbial net-works. FEMS microbiology reviews. 2018;42(6):761–780.

[45] Novitsky V, Steingrimsson JA, Howison M, Gillani FS, Li Y, Manne A, et al. Empirical comparison of analytical approaches for identifying molecular HIV-1 clusters. Scientific reports. 2020;10(1):1–11.

[46] Volz EM, Koopman JS, Ward MJ, Brown AL, Frost SD. Simple epidemiological dynamics explain phylogenetic clustering of HIV from patients with recent infection. PLoS Comput Biol. 2012;8(6):e1002552.

[47] Novitsky V, Moyo S, Lei Q, DeGruttola V, Essex M. Impact of sampling density on the extent of HIV clustering. AIDS research and human retroviruses. 2014;30(12):1226–1235.

[48] Chato C, Kalish ML, Poon AF. Public health in genetic spaces: a statistical framework to optimize cluster-based outbreak detection. Virus evolution. 2020;6(1):veaa011.

[49] Matsen FA, Kodner RB, Armbrust EV. pplacer: linear time maximum-likelihood and Bayesian phylogenetic placement of sequences onto a fixed reference tree. BMC bioinformatics. 2010;11(1):538.

[50] Dennis AM, Volz E, Frost AMSD, Hossain M, Poon AF, Rebeiro PF, et al. HIV-1 transmission clustering and phylodynamics highlight the important role of young men who have sex with men. AIDS research and human retroviruses. 2018;34(10):879–888.

[51] Wolf E, Herbeck JT, Van Rompaey S, Kitahata M, Thomas K, Pepper G, et al. Phylogenetic evidence of HIV-1 transmission between adult and adolescent men who have sex with men. AIDS research and human retroviruses. 2017;33(4):318–322.

[52] Vrancken B, Adachi D, Benedet M, Singh A, Read R, Shafran S, et al. The multi-faceted dynamics of HIV-1 transmission in Northern Alberta: A combined analysis of virus genetic and public health data. Infection, Genetics and Evolution. 2017;52:100–105.

[53] Ye J, Hao M, Xing H, Zhang F, Wu H, Lv W, et al. Transmitted HIV drug resistance among individuals with newly diagnosed HIV infection: a multicenter observational study. Aids. 2020;34(4):609–619.

[54] Tordoff D, Herbeck J, Buskin S, Lechtenberg R, Golden M, Kerani R. O19.4 Molecular epidemiology of HIV among foreign-born residents of King County, Washington, USA, using HIV surveillance data. BMJ. 2019;95(Suppl 1):A83.

[55] Kosakovsky Pond SL, Posada D, Stawiski E, Chappey C, Poon AF, Hughes G, et al. An evolutionary model-based algorithm for accurate phylogenetic breakpoint mapping and subtype prediction in HIV-1. PLoS computational biology. 2009;5(11):e1000581.

[56] Nguyen LT, Schmidt HA, Von Haeseler A, Minh BQ. IQ-TREE: a fast and effective stochastic algorithm for estimating maximum-likelihood phylogenies. Molecular biology and evolution. 2015;32(1):268–274.

[57] Price MN, Dehal PS, Arkin AP. FastTree 2–approximately maximum-likelihood trees for large alignments. PloS one. 2010;5(3):e9490.

[58] Revell LJ. phytools: an R package for phylogenetic comparative biology (and other things). Methods in ecology and evolution. 2012;3(2):217–223.

[59] Paradis E, Schliep K. ape 5.0: an environment for modern phylogenetics and evolutionary analyses in R. Bioinformatics. 2019;35(3):526–528.

[60] Matsen IV FA, Evans SN. Edge principal components and squash clustering: using the special structure of phylogenetic placement data for sample comparison. PloS one. 2013;8(3):e56859.

[61] Akaike H. Information theory and an extension of the maximum likelihood principle. In: Selected papers of hirotugu akaike. Springer; 1998. p. 199–213.

[62] Robin X, Turck N, Hainard A, Tiberti N, Lisacek F, Sanchez JC, et al. pROC: an open-source package for R and S+ to analyze and compare ROC curves. BMC bioinformatics. 2011;12(1):1–8.

[63] Vrancken B, Rambaut A, Suchard MA, Drummond A, Baele G, Derdelinckx I, et al. The genealogical population dynamics of HIV-1 in a large transmission chain: bridging within and among host evolutionary rates. PLoS Comput Biol. 2014;10(4):e1003505.

[64] Robertson DL, Anderson J, Bradac J, Carr J, Foley B, Funkhouser R, et al. HIV-1 nomenclature proposal. Science. 2000;288(5463):55–55.

[65] Stimson J, Gardy J, Mathema B, Crudu V, Cohen T, Colijn C. Beyond the SNP threshold: identifying outbreak clusters using inferred transmissions. Molecular biology and evolution. 2019;36(3):587–603.

[66] Swift A, Liu L, Uber J. Reducing MAUP bias of correlation statistics between water quality and GI illness. Computers, Environment and Urban Systems. 2008;32(2):134–148.

[67] Nakaya T, Fotheringham AS, Brunsdon C, Charlton M. Geographically weighted Poisson regression for disease association mapping. Statistics in medicine. 2005;24(17):2695–2717.

[68] Nakaya T. An information statistical approach to the modifiable areal unit problem in incidence rate maps. Environment and Planning A. 2000;32(1):91–109.

[69] Bortz D, Nelson P. Model selection and mixed-effects modeling of HIV infection dynamics. Bulletin of mathematical biology. 2006;68(8):2005–2025.

[70] Rentsch C, Bebu I, Guest JL, Rimland D, Agan BK, Marconi V. Combining epidemiologic and biostatistical tools to enhance variable selection in HIV cohort analyses. PloS one. 2014;9(1):e87352.

[71] Novitsky V, Moyo S, Lei Q, DeGruttola V, Essex M. Importance of viral sequence length and number of variable and informative sites in analysis of HIV clustering. AIDS research and human retroviruses. 2015;31(5):531–542.

[72] Coltart CE, Hoppe A, Parker M, Dawson L, Amon JJ, Simwinga M, et al. Ethical considerations in global HIV phylogenetic research. The lancet HIV. 2018;5(11):e656–e666.

[73] Le Vu S, Ratmann O, Delpech V, Brown AE, Gill ON, Tostevin A, et al. Comparison of cluster-based and source-attribution methods for estimating transmission risk using large HIV sequence databases. Epidemics. 2018;23:1–10.

[74] Bachmann N, Kusejko K, Nguyen H, Chaudron SE, Kadelka C, Turk T, et al. Phylogenetic Cluster Analysis Identifies Virological and Behavioral Drivers of HIV Transmission in MSM. Clinical Infectious Diseases. 2020;72(12):2175–2183.

[75] Morel B, Barbera P, Czech L, Bettisworth B, Hübner L, Lutteropp S, et al. Phylogenetic anal-ysis of SARS-CoV-2 data is difficult. Molecular biology and evolution. 2021;38(5):1777–1791.

